# Anatomical mapping of dopamine D1 and muscarinic M4 receptor co-expression

**DOI:** 10.64898/2026.09.27.754786

**Authors:** Rian Garland, Maya Dickson, Alice Lee, Ali Noorouzi, Hayato Umaoka, Ana Semeano, James R. Monaghan, Hideaki Yano

**Affiliations:** Department of Pharmaceutical and Biomedical Sciences, Center for Drug Discovery, School of Pharmacy and Pharmaceutical Sciences, Bouvé College of Health Sciences Northeastern University, Boston, Massachusetts 02115, United States; Department of Biology, Northeastern University, Boston, Massachusetts 02115, United States

**Keywords:** dopamine D1 receptor, muscarinic M4 receptor, striatum, nucleus accumbens, reporter mouse, HCR-FISH

## Abstract

Dopamine D1 receptors (D1R) and muscarinic M4 receptors (M4R) regulate overlapping functions in the cortex and the striatum, but the cellular relationship between D1R- and M4R-associated populations across brain regions remains incompletely defined. Using D1R-tdTomato/M4R-EGFP double-reporter mice, we quantified tdTomato-only, EGFP-only, and coexpressing cellular populations across the dorsal striatum, nucleus accumbens, and medial prefrontal cortex in 18 animals. Reporter composition differed strongly by anatomical region. Coexpression was prominent throughout the dorsal striatum and greatest in the dorsomedial striatum, whereas the nucleus accumbens shifted toward greater tdTomato-only and reduced EGFP-only representation, particularly in the core. In contrast, the medial prefrontal cortex was dominated by EGFP-only cells. These regional patterns were broadly retained across sex and age. HCR-FISH further supported spatial correspondence between endogenous *Drd1* and *Chrm4* transcript signal and the respective fluorescent reporters. Together, these findings demonstrate region-dependent organization of D1R- and M4R-associated cellular populations across the cortex and striatum and define distinct anatomical contexts in which M4R-directed signaling may influence D1R-associated neurons.

## Introduction

Dopamine D1 receptor (D1R) signaling is a major component of basal ganglia function and is particularly important within direct-pathway striatal projection neurons, where D1R activation promotes Gαs/olf-dependent stimulation of adenylyl cyclase and downstream cAMP signaling.^1–3^ These pathways contribute to the regulation of neuronal excitability, synaptic transmission, and longer-term cellular responses, but the functional consequences of D1R activation are shaped by the receptor and signaling systems expressed within the same neurons.^2,4–6^ Among these, the muscarinic M4 receptor (M4R) is especially relevant because it couples primarily to Gαi/o proteins and can inhibit adenylyl cyclase activity, reduce intracellular cAMP, and regulate neurotransmitter release.^7–10^ At the level of cAMP production, M4R is therefore positioned to counterbalance the canonical stimulatory D1R-Gαs/olf pathway within cells that express both receptors. More broadly, dopaminergic and cholinergic signaling are tightly intertwined within the striatum, where changes in their relative influence can substantially alter circuit function and behavior.^7,11–16^

This relationship has an established cellular and functional basis in the striatum. D1R and M4R have been reported to be coexpressed in a subset of striatonigral medium spiny neurons,^17^ creating the opportunity for dopaminergic and cholinergic signals to converge within the same neuronal population. M4R is also the functionally predominant muscarinic receptor subtype in mouse and rat striatum,^18^ further supporting its importance in shaping local dopamine-responsive signaling. Consistent with this role, selective deletion of M4R from D1R-expressing neurons disrupts muscarinic inhibition of D1R-stimulated cAMP signaling and enhances dopamine-dependent locomotor and psychostimulant responses.^19^ Together, these findings establish M4R as a functionally important regulator within D1R-associated neuronal populations and provide a strong rationale for defining where these two receptor systems are anatomically positioned to interact.

The capacity of M4R to regulate dopamine-responsive signaling has also made the receptor an important pharmacological target. Selective activation or positive allosteric modulation of M4R produces antipsychotic-like effects in preclinical models and can attenuate behavioral and neurochemical responses to psychostimulants, including amphetamine and cocaine.^20–25^ M4R signaling has additionally been implicated in the regulation of corticostriatal plasticity, where enhancement of M4R function can oppose D1R-dependent potentiation and ameliorate abnormal plasticity associated with L-DOPA-induced dyskinesia.^26^ These findings suggest that M4R-directed pharmacology can influence dopamine-dependent circuit function without requiring direct pharmacological manipulation of D1R itself.

Interest in this strategy has extended into clinical drug development. The efficacy of xanomeline–trospium, which combines an M1R/M4R-preferring central muscarinic agonist with a peripherally restricted muscarinic antagonist, provides clinical evidence that therapeutically useful modulation of central muscarinic signaling is achievable.^27–29^ The parallel development of compounds designed to more selectively engage M4R further reflects continued interest in separating the contribution of this receptor from broader muscarinic activation.^30,31^ However, molecular selectivity for M4R does not by itself determine which dopamine-responsive neuronal populations will be affected. The functional reach of an M4R-directed intervention will also depend on the regional and cellular distribution of M4R relative to D1R-associated populations. Defining that distribution is therefore relevant not only to understanding D1R–M4R biology, but also to evaluating the circuit selectivity that M4R-targeted pharmacology may realistically provide.

The potential consequences of D1R–M4R coexpression are also likely to depend strongly on anatomical context. D1R signaling contributes to distinct functions across the regions examined here, and even within the striatum, neighboring subdivisions participate in different behavioral and circuit processes. In the dorsal striatum, D1R-expressing direct-pathway neurons contribute to basal ganglia output related to movement and action selection.^32–36^ The dorsolateral striatum is more strongly associated with sensorimotor processing and habit-related control, whereas the dorsomedial striatum contributes more prominently to associative and goal-directed behavior.^37,38^ The nucleus accumbens extends this comparison into ventral striatal circuitry involved in motivation, reinforcement, and reward-related behavior.^34–36^ Its core and shell differ in connectivity, dopaminergic innervation, and contributions to motivated behavior, with the core more strongly associated with cue-guided reward seeking and the shell with contextual regulation of reward-related behavior.^39–43^

Extra-striatal cortex provides an important contrast to this established striatal relationship. D1R signaling in the medial prefrontal cortex contributes to working memory, executive function, and adaptive behavior,^44–47^ and previous conditional deletion of M4R from D1R-expressing cells produced marked reductions in M4R expression in the caudate-putamen and nucleus accumbens but not in frontal cortex.^24^ This finding suggests that cortical M4R expression may be distributed substantially among cellular populations that do not express D1R, providing a specific reason to compare cortical and striatal regions rather than assuming that the extensive D1R-M4R association described in striatal neurons generalizes throughout the forebrain. Examining dorsal and ventral striatal and medial prefrontal cortical regions within the same experimental framework can therefore test whether D1R- and M4R-associated populations retain a common cellular organization or instead exhibit pronounced regional specialization.

Age and sex may represent additional sources of variation in dopamine-associated circuitry. The prevalence and clinical presentation of disorders involving dopaminergic systems vary across age and sex,^48–50^ and evoked dopamine release in the dorsal striatum has likewise been reported to differ across these biological variables.^51^ Whether age or sex also influences the regional cellular relationship between D1R- and M4R-associated populations remains unclear. Examining these variables alongside regional organization can therefore help determine whether broader patterns of D1R–M4R association are stable across the experimental population or vary with biological context. Despite evidence for D1R-M4R coexpression in striatal neurons, the cellular organization of these receptor-associated populations across regions remains incompletely resolved. D1R and M4R have been identified within the same striatonigral neurons,^17^ and selective deletion of M4R from D1R-expressing cells produces markedly different effects on M4R expression in striatal versus cortical regions.^24^ However, these observations do not establish the relative proportions of D1R-associated, M4R-associated, and coexpressing cells within each region or determine how those proportions vary across anatomically distinct circuits. This distinction is important because detection of both receptors within a brain region can arise from extensive same-cell coexpression, largely separate cellular populations, or intermediate combinations of the two. Each arrangement would define a different cellular opportunity for M4R to influence D1R-associated neurons.

Direct assessment of this organization is further complicated by the challenges of protein-level localization for G protein-coupled receptors, including limitations in the specificity and reliability of available antibodies.^52^ Reporter-based approaches therefore provide a practical means of resolving D1R- and M4R-associated populations within the same tissue, while transcript-level measurements can provide complementary evidence that reporter-defined populations correspond to endogenous receptor expression. A direct regional comparison of single-positive and coexpressing populations is needed to determine whether the strong D1R-M4R association established in striatal neurons is retained across dorsal and ventral striatal compartments and whether a different cellular organization characterizes medial prefrontal cortex. Defining these patterns provides the anatomical basis for interpreting where M4R-directed pharmacology is positioned to act within D1R-associated neuronal populations.

To our knowledge, no previous study has systematically compared the relative representation of D1R-associated, M4R-associated, and coexpressing cellular populations across dorsal striatal, accumbal, and medial prefrontal cortical regions within the same experimental framework. Here, we asked whether the cellular relationship between D1R- and M4R-associated populations is conserved across regions or instead reflects the anatomical and functional specialization of distinct forebrain circuits, and whether these broader patterns are retained across age and sex. Using D1R-tdTomato/M4R-EGFP double-reporter mice, we compared the relative representation of D1R-associated, M4R-associated, and coexpressing populations across the dorsolateral and dorsomedial striatum, nucleus accumbens core and shell, and medial prefrontal cortex. Reporter-based mapping was complemented by HCR-FISH to assess correspondence with endogenous *Drd1* and *Chrm4* transcripts. By examining these populations within a common anatomical framework, we sought to define where D1R and M4R occupy shared cellular populations, where their distributions diverge, and whether these relationships remain broadly consistent across age and sex. Defining this regional organization provides a cellular framework for interpreting the potential reach and selectivity of M4R-mediated modulation within D1R-associated neuronal systems.

## Methods

### Animals, ethics, and reporter mouse model

D1R- and M4R-associated cell populations were examined in double-reporter mice generated by breeding B6.Cg-Tg(*Drd1a*-tdTomato)6Calak/J (D1Tomato) mice with Tg(*Chrm4*-EGFP)Y86Gsat (M4EGFP) mice. Both parental strains are bacterial artificial chromosome transgenic reporter lines. In the resulting tissue, tdTomato fluorescence marks the D1R-associated reporter population, whereas EGFP fluorescence marks the M4R-associated reporter population. Only offspring carrying both reporter transgenes were included in the coexpression analyses. (**Fig. 1A-C**).

**Figure 1.**
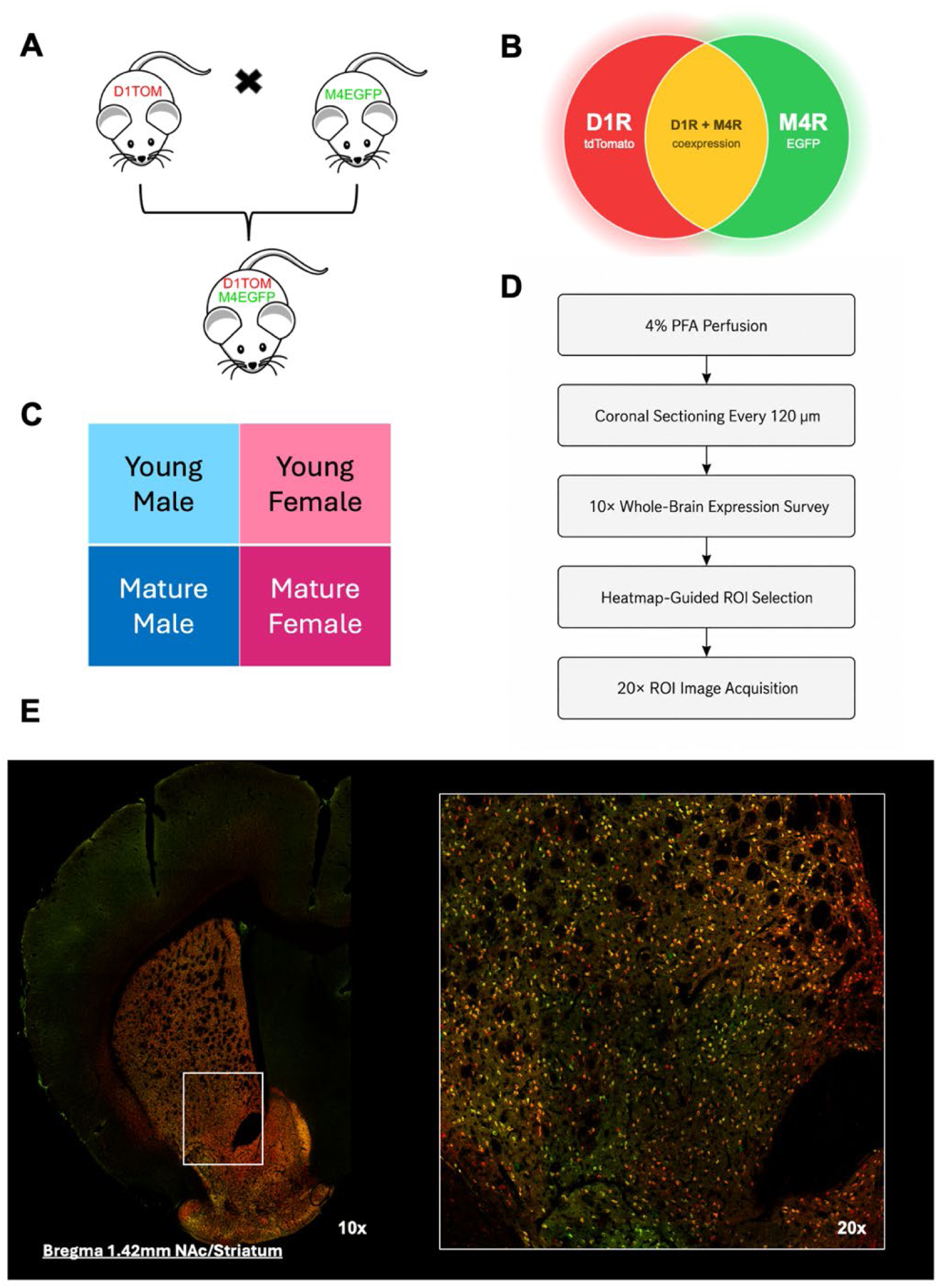
Double-reporter model and quantitative imaging workflow for analysis of D1R- and M4R-associated reporter populations. **A.** D1R-tdTomato and M4R-EGFP single-reporter mice were crossed to generate D1R-tdTomato/M4R-EGFP double-reporter animals. **B.** Schematic representation of D1R+ (tdTomato-only), M4R+ (EGFP-only), and D1R+/M4R+ coexpressing reporter populations. **C.** Experimental groups included young male, young female, mature male, and mature female animals. Young animals were 8–12 weeks of age and mature animals were 22–28 weeks of age. **D.** Tissue and imaging workflow consisting of 4% paraformaldehyde perfusion, coronal section sampling at 120 µm intervals, 10× whole-brain survey imaging, heatmap-guided region-of-interest selection, and 20× ROI image acquisition. **E.** Representative 10× coronal image and corresponding 20× enlargement from the nucleus accumbens/striatal region. Red indicates D1R-associated tdTomato fluorescence, green indicates M4R-associated EGFP fluorescence, and yellow indicates overlapping reporter signal.

Male and female mice from two age groups were studied: young adults aged 8–12 weeks and mature adults aged 22–28 weeks. Genotyping was performed by Transnetyx using tail-clip samples to detect the *Drd1a*-tdTomato and *Chrm4*-EGFP transgenes. Tissue from 18 double-positive mice was included in the study.

All procedures involving animals were approved by the Northeastern University Institutional Animal Care and Use Committee under protocol 25-0923R and were conducted at 140 The Fenway under Animal Welfare Assurance D16-00095. Animal care and experimental procedures followed the NIH Guide for the Care and Use of Laboratory Animals. Mice had ad libitum access to food and water and were routinely monitored for signs of illness, pain, or distress.

### Perfusion, fixation, cryoprotection, and sectioning

Mice were anesthetized with isoflurane and transcardially perfused with 30 mL ice-cold phosphate-buffered saline (PBS), followed by 30 mL 4% paraformaldehyde (PFA). Brains were removed and immersed in approximately 10 mL 4% PFA at 4°C overnight for post-fixation for no longer than 24 h. Following post-fixation, tissue was either transferred directly to sucrose for cryoprotection or maintained in PBS at 4°C until further processing. (**Fig. 1D**).

Brains were cryoprotected sequentially in 15% and 30% (w/v) sucrose at 4°C for approximately 24 h in each solution, with fresh sucrose solution provided after approximately 12 h. After cryoprotection, excess solution was removed with lint-free tissue, and brains were embedded in optimal cutting temperature compound and frozen at −20°C for at least 1–2 h. Embedded tissue was stored at −20°C or −80°C until sectioning.

Coronal sections were cut at 40 µm using either a Microm HM 550 or Leica CM1860 cryostat. Approximately every third section was collected directly onto Superfrost Plus microscope slides, corresponding to a nominal 120-µm interval between collected sections. Slide-mounted sections were stored at −20°C until further processing.

### Reporter tissue preparation

Slide-mounted sections designated for direct reporter fluorescence imaging were removed from −20°C storage and allowed to equilibrate to room temperature. Sections were then washed in phosphate-buffered saline (PBS) for approximately 15 min. Excess buffer was gently removed, and the tissue was lightly dried before coverslipping with ProLong Diamond Antifade Mountant containing DAPI. Mounted slides were allowed to cure for at least overnight before the coverslip edges were sealed with clear nail polish. Slides were stored at 4°C protected from light until imaging.

### HCR-FISH reporter validation

Hybridization chain reaction fluorescence in situ hybridization (HCR-FISH) was performed on slide-mounted sections to assess correspondence between endogenous receptor transcripts and reporter fluorescence. Separate Molecular Instruments probe sets targeting *Drd1* and *Chrm4* were used because both target probe sets were paired with the B3 amplifier and Alexa Fluor 647-conjugated hairpins; the two transcripts were therefore evaluated on separate tissue sections rather than multiplexed within the same section. A *dapB* negative-control probe was processed in parallel on separate sections for comparison with each reporter dataset. Validation tissue was obtained from one double-reporter animal; at least two sections were analyzed for each target probe, and one section was analyzed for the *dapB* negative control.

HCR-FISH was performed using an established V3 HCR-FISH protocol. Sections were washed three times for 5 min in 1× phosphate-buffered saline (PBS) at room temperature and incubated overnight in 70% ethanol as the tissue pretreatment step. Proteinase K digestion was not used. Sections were then prehybridized in 30% hybridization buffer for 10 min at 37°C. Probe stocks (1 µM) were diluted 1:200 in 30% hybridization buffer, and sufficient probe solution was applied to cover each tissue section. Sections were covered with parafilm and incubated overnight at 37°C in a humidified chamber.

Following hybridization, sections were washed three times for 15 min at 37°C in 30% probe wash buffer and twice for 5 min in 5× saline-sodium citrate containing Tween-20 (SSCT) at room temperature. Sections were subsequently incubated in amplification buffer for 30 min at room temperature. Alexa Fluor 647-conjugated B3 hairpins were heated to 95°C for 90 s and allowed to cool to room temperature for at least 30 min in the dark before use. Hairpins were diluted 1:50 in amplification buffer, and less than 50 µL was applied to each section. Sections were covered with parafilm and incubated overnight at room temperature in a humidified chamber protected from light.

After amplification, sections were washed twice for 15 min in 5× SSCT at room temperature. Nuclei were counterstained with DAPI for 5 min, followed by a 5-min PBS wash. Sections were coverslipped with ProLong Diamond Antifade Mountant without DAPI, sealed with clear nail polish, and stored at 4°C protected from light until imaging.

### Confocal image acquisition

Sections were imaged using an Olympus FV3000 confocal laser-scanning microscope. A 10× objective was used for anatomical surveys and regional orientation, while images used for cell-level quantification were acquired with a 20× objective at 1× optical zoom. Quantitative image stacks were collected at 2048 × 2048 pixels, with five optical planes spanning approximately 20 µm. Maximum-intensity projections were generated for subsequent image analysis. Imaging parameters were kept constant across all samples included in quantitative comparisons. (**Fig. 1E**).

Reporter fluorescence was acquired in two scan phases. DAPI and tdTomato were imaged together using 405- and 561-nm excitation, respectively. DAPI emission was collected from 430–470 nm at 1% laser power and 750 V, and tdTomato emission from 570–670 nm at 1% laser power and 800 V. EGFP was acquired separately using 488-nm excitation, with emission collected from 500–600 nm at 1% laser power and 700 V. Images were acquired by line-sequential scanning without signal averaging. Detector gain was set to 1.000, offset to 0%, and a 10% laser neutral-density filter was used. The confocal aperture was set automatically by the acquisition software.

HCR-FISH sections were imaged with a 60× oil-immersion objective at 1× optical zoom. DAPI, tdTomato, EGFP, and Alexa Fluor 647 were excited at 405, 561, 488, and 640 nm, with emission collected at 425–475, 575–625, 500–550, and 650–750 nm, respectively. Z-stacks consisted of 11 optical planes spanning approximately 5 µm. The 640-nm laser was typically set to 5% power, with detector voltage adjusted to resolve punctate HCR signal while minimizing background. Maximum-intensity projections were generated for visualization and puncta analysis.

Original Olympus .oir files and associated metadata were retained as source data, and all derived images used for visualization or quantitative analysis were generated from copies of the original files.

### 10× heatmap generation and ROI prioritization

Half-brain images acquired at 10× magnification were used to generate threshold-based reporter heatmaps with a custom Python workflow. RGB composites containing the tdTomato and EGFP channels were converted to 8-bit grayscale using luminance weighting according to 0.299R + 0.587G + 0.114B. For each image, mean grayscale intensity was calculated after excluding pixels with intensity values of 0 or 255. These image-level means were then averaged across the full comparison set to establish a common reference value. (**Fig. 2**).

**Figure 2.**
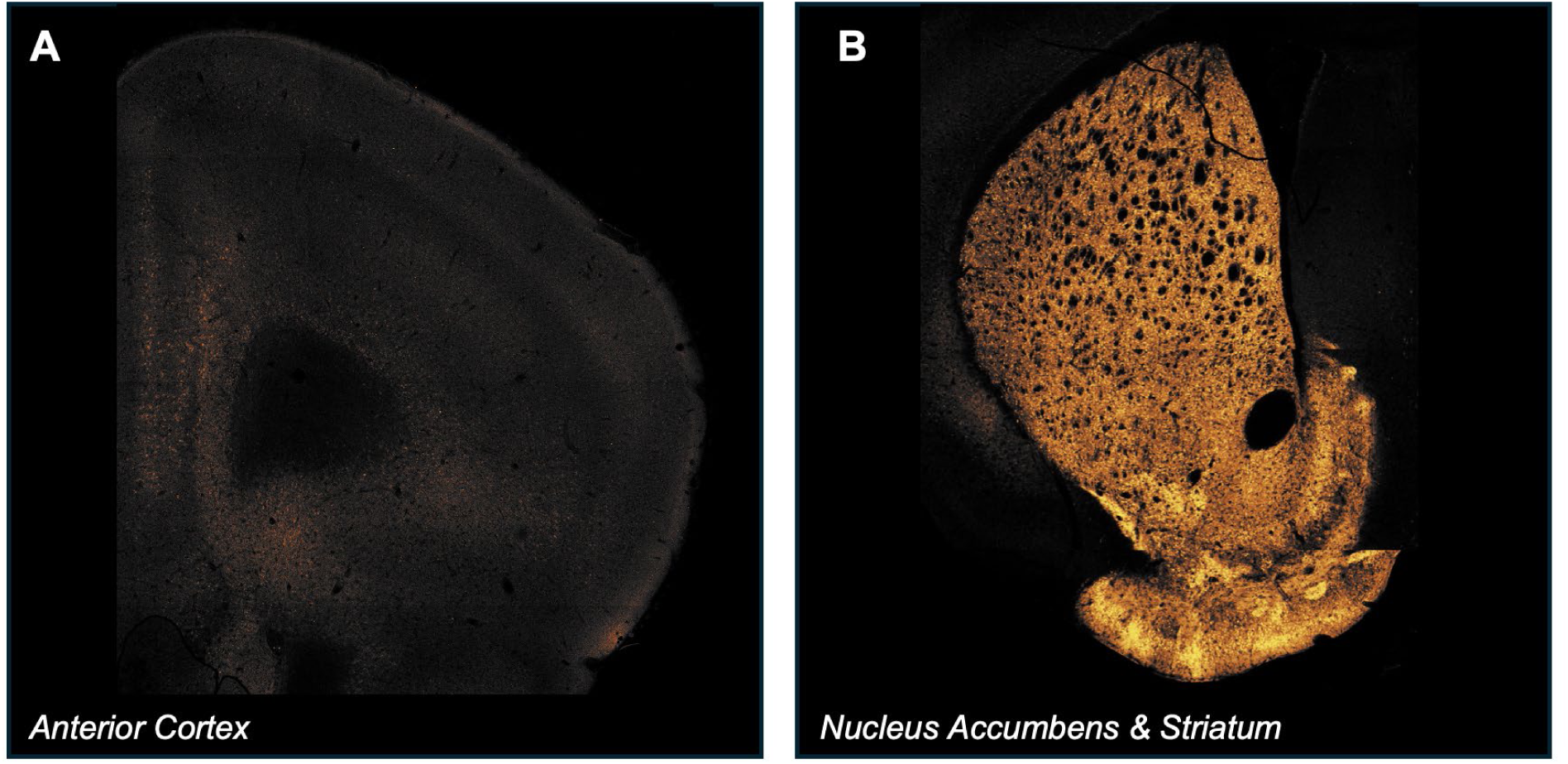
Representative 10× survey heatmaps used to guide regional sampling. **A.** Representative anterior cortical section showing areas of comparatively high combined tdTomato and EGFP reporter signal. **B.** Representative nucleus accumbens and striatal section showing prominent reporter signal across dorsal and ventral striatal regions. Heatmaps were generated from the combined reporter channels and used to guide subsequent 20× region-of-interest selection. Heatmap intensity was used for anatomical survey and field selection and was not interpreted as a quantitative comparison of reporter expression between regions or animals.

The heatmap threshold was set at three times this overall mean, corresponding to 300% of the comparison-set average. Pixels meeting or exceeding this threshold were displayed with the autumn colormap at 0.6 opacity over the underlying grayscale image, and the same absolute threshold was used for all images within a comparison set. Heatmaps were used to identify areas of comparatively high combined reporter signal and to guide selection of regions for higher-magnification imaging. They were not used to classify individual cells or to make quantitative comparisons of fluorescence intensity between animals.

### Section selection, atlas alignment, and ROI definitions

Anatomical regions were identified within the serial whole-brain section set by comparison with Paxinos and Franklin’s The Mouse Brain in Stereotaxic Coordinates. Regional morphology, recognizable anatomical landmarks, and rostrocaudal progression through the section series were considered together when assigning regions. Sampling for a given structure began at the first mounted section in which its boundaries could be identified reliably and continued at approximately every third mounted section while the structure remained present. Sampling intervals could vary when tissue quality or anatomical boundaries prevented reliable region identification. Individual sections were therefore not assigned formal bregma coordinates or atlas plate numbers.

Because the rectangular confocal field frequently extended beyond the anatomical structure of interest, regions of interest (ROIs) were manually delineated in Fiji before quantitative analysis. Areas outside the intended anatomical boundaries, non-tissue spaces, fiber-tract openings, and damaged or missing tissue were excluded. For the dorsal striatum, the dorsal third of the structure was selected and divided mediolaterally into dorsolateral (DLSTR) and dorsomedial (DMSTR) ROIs. The nucleus accumbens (NAc) core was defined as the compact region surrounding the anterior commissure, whereas the NAc shell comprised the more medial and peripheral accumbal tissue surrounding the core and lying dorsal to the olfactory tubercle. The medial prefrontal cortex (mPFC) ROI included anterior cingulate, prelimbic, and infralimbic cortex. These cortical subdivisions were analyzed together because their internal borders could not be identified consistently at the imaging scale used. (**Fig. 3**).

**Figure 3.**
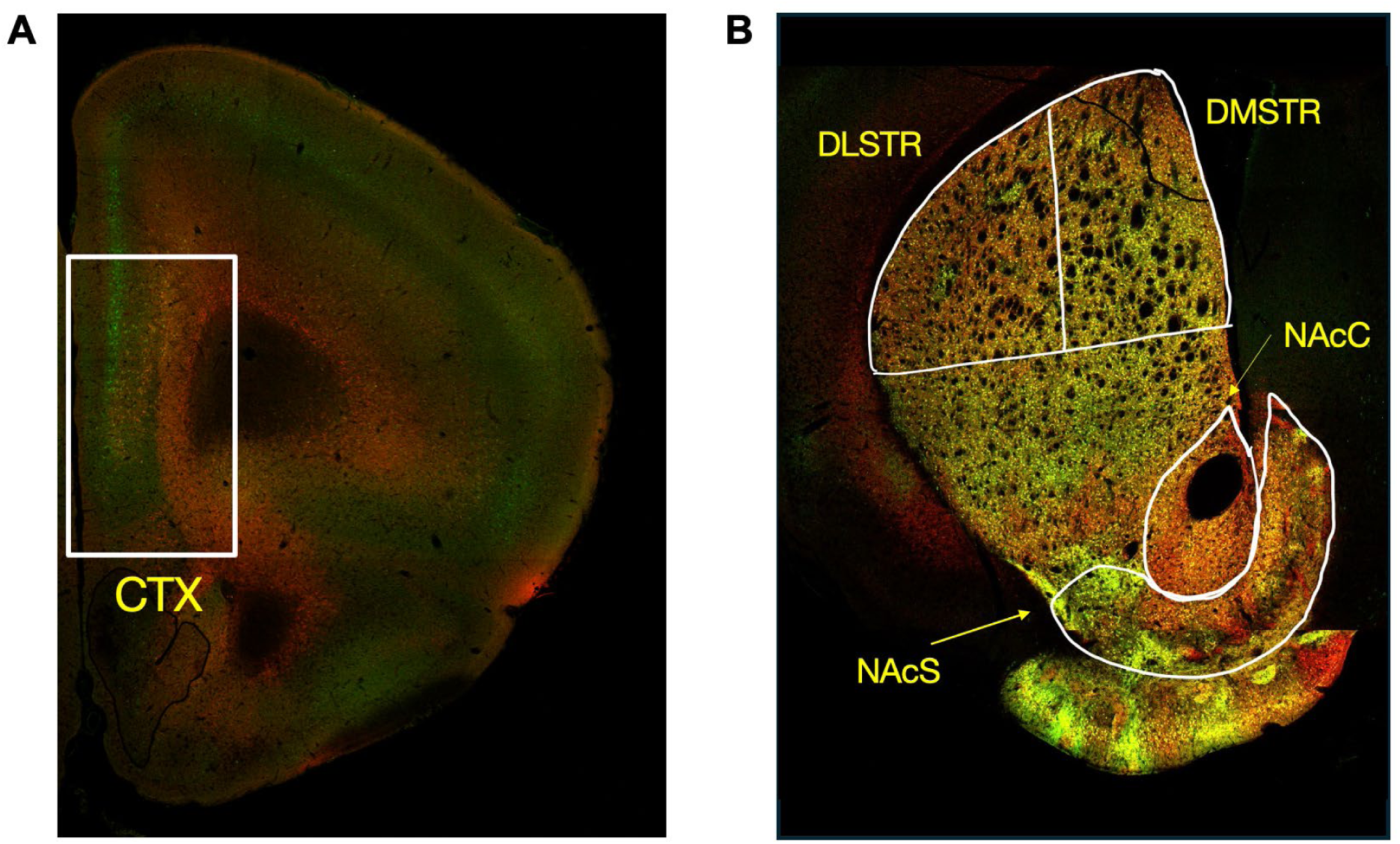
Anatomical definition of quantified cortical and striatal regions of interest. **A.** Representative coronal section showing the sampled medial prefrontal cortex region of interest. **B.** Representative coronal section showing operational boundaries for the dorsolateral striatum (DLSTR), dorsomedial striatum (DMSTR), nucleus accumbens core (NAcC), and nucleus accumbens shell (NAcS). White outlines indicate regions used for subsequent quantitative analysis.

The right hemisphere was preferentially used for imaging and quantification. When folding, tissue damage, incomplete mounting, or poorly preserved anatomical boundaries compromised the right hemisphere, the corresponding left-hemisphere region was used instead; hemisphere was not treated as an experimental variable. Sections or fields were excluded when tissue damage, mounting defects, acquisition artifacts, or inadequate anatomical correspondence prevented reliable ROI delineation. ROI selection and image analysis were not formally blinded to animal identity or experimental group, although group metadata were not used to guide anatomical boundary placement, field selection, or exclusion decisions. At least three sections were analyzed for each animal-region combination. Section-level counts were treated as subsamples and aggregated within animal before reporter-class composition was calculated.

### 20× image preprocessing and soma classification

Images acquired at 20× were processed separately from the 10× heatmap workflow. Five-plane z-stacks were converted to maximum-intensity projections while preserving the DAPI, tdTomato, and EGFP channels separately. Anatomical ROIs were manually traced on the projected images in Fiji and saved in the ROI Manager. To prepare images for automated analysis, each selected ROI was applied to a duplicate of the projected hyperstack, signal outside the ROI boundary was removed, and the remaining image was cropped to the rectangular bounds of the ROI. Cropped ROIs were exported as three-page TIFF files containing DAPI, tdTomato, and EGFP channels in a consistent order. Original images and uncropped projections were retained separately.

Cell detection and reporter classification were performed using a custom Python pipeline. No additional contrast enhancement, denoising, background subtraction, or morphological preprocessing was applied before segmentation. A binary DAPI mask was generated using a fixed intensity threshold of 200. The tdTomato and EGFP channels were segmented independently with the pretrained Cellpose cyto3 model, using automatic object-diameter estimation, a flow threshold of 0.4, a cell-probability threshold of 0.0, and augmentation enabled. Reporter objects were retained only when their contours overlapped at least five positive pixels in the DAPI mask, thereby restricting classification to reporter signal associated with nuclear staining. (**Fig. 4**).

**Figure 4.**
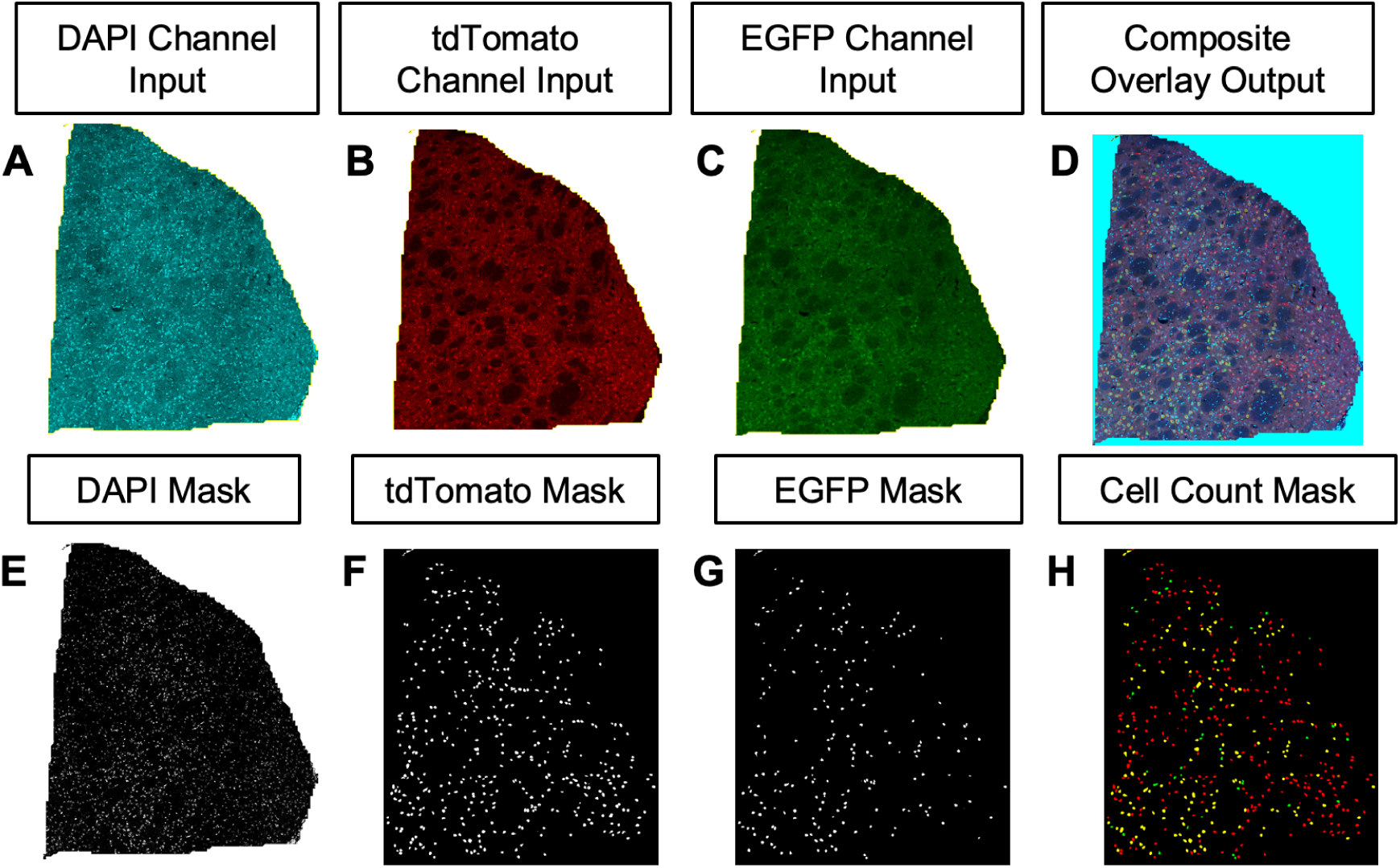
DAPI-gated image-analysis pipeline for reporter-defined cell population classification. **A.** DAPI channel input. **B.** tdTomato channel input. **C.** EGFP channel input. **D.** Composite overlay image. **E.** DAPI mask used to retain reporter-associated objects corresponding to nucleated cells. **F.** tdTomato mask. **G.** EGFP mask. **H.** Final classified-cell output used to assign tdTomato-only, EGFP-only, and coexpressing cell populations.

DAPI-gated tdTomato and EGFP objects were compared spatially to assign reporter category. A tdTomato contour and an EGFP contour were classified as coexpressing when their intersection-over-union value was at least 0.35; matched contours were combined into a single coexpressing object, while unmatched contours were retained as tdTomato-only or EGFP-only objects. The pipeline therefore quantified tdTomato-only, EGFP-only, and coexpressing reporter populations but did not enumerate reporter-negative DAPI-only cells. The same analysis parameters were applied throughout the comparison set.

During pipeline development, segmentation masks and classification overlays were visually compared with the corresponding fluorescence images, and automated counts were checked against informal manual counts in representative fields to identify obvious segmentation or classification errors. These checks informed parameter selection but were not treated as a formal validation dataset. The analysis was implemented in Python 3.12.9 using Cellpose 3.1.1.1.

### Automated HCR-FISH puncta analysis

Automated image analysis was performed in Python to quantify spatial association between HCR-FISH puncta and the corresponding reporter fluorescence. For *Chrm4* sections, Alexa Fluor 647 puncta were evaluated relative to M4-EGFP fluorescence; for *Drd1* sections, Alexa Fluor 647 puncta were evaluated relative to tdTomato fluorescence. DAPI was not included in the puncta-overlap analysis.

For both probe sets, slowly varying background in the far-red channel was reduced by subtracting a Gaussian-blurred copy of the image generated with a sigma of 4.0 µm, followed by Gaussian smoothing with a sigma of 0.15 µm. For target-probe images, far-red puncta were detected using a threshold defined as the background median plus seven robust standard deviations. Because the four-channel *dapB* negative-control dataset exhibited higher far-red background, puncta in that dataset were detected using the background median plus nine robust standard deviations. Connected objects with estimated diameters between 0.5 and 10 µm were retained as HCR-FISH puncta.

The corresponding reporter channel was converted to a binary mask. For target-probe images, the reporter-positive threshold was set to the median reporter intensity within each analyzed field. For the four-channel *dapB* negative-control dataset, reporter thresholds were predefined slightly above the corresponding target-condition thresholds to account for the different acquisition and background characteristics. To accommodate minor spatial displacement between fluorescence channels, the sampling footprint surrounding each detected HCR-FISH punctum was expanded by 0.2 µm before reporter overlap was measured. A punctum was classified as reporter-associated when at least 10% of its sampling footprint overlapped the reporter-positive mask. Punctum area, mean far-red intensity, mean reporter intensity, fraction of reporter overlap, and overlap classification were recorded. Quality-control images were reviewed to verify punctum detection, reporter-overlap classification, and threshold performance before quantitative interpretation.

### Data export, aggregation, and statistical analysis

The image-analysis pipeline generated counts for three mutually exclusive reporter-defined populations: tdTomato-only, EGFP-only, and tdTomato-EGFP coexpressing cells. Counts from all included images and sections were aggregated by animal and anatomical region before tissue-level outcomes were calculated. Images, ROIs, and sections were treated as subsamples rather than independent biological replicates, with the individual animal serving as the experimental unit.

For each animal-region combination, the total reporter-classified population was calculated as the sum of the tdTomato-only, EGFP-only, and coexpressing populations. Reporter composition was expressed as the percentage of this total represented by each category. Because reporter-negative DAPI-only nuclei were not enumerated by the analysis pipeline, these percentages describe the composition of the reporter-classified population rather than the percentage of all cells within an ROI. Absolute category counts were retained as measures of sampling yield but were not normalized to ROI area or interpreted as estimates of cell density.

Statistical analyses were performed using GraphPad Prism 10. Reporter-defined population composition across the dorsolateral striatum, dorsomedial striatum, nucleus accumbens core, and nucleus accumbens shell was analyzed in the pooled cohort (n = 18) using a two-way repeated-measures ANOVA with anatomical region and reporter category as within-animal factors. Geisser-Greenhouse correction was applied to account for potential violations of sphericity. Tukey’s multiple-comparisons test was used to compare reporter categories within anatomical regions and anatomical regions within reporter categories, with reported post hoc p values adjusted for multiple comparisons. To provide reporter-specific omnibus tests for the regional comparisons, tdTomato-only,

EGFP-only, and coexpressing percentages were also analyzed separately across the four striatal regions using one-way repeated-measures ANOVA with Geisser-Greenhouse correction and Tukey-adjusted pairwise comparisons.

The same striatal analyses were repeated after stratification by sex (female, n = 9; male, n = 9) and age (young, n = 9; mature, n = 9). Combined age-by-sex groups were also analyzed using the same reporter-specific regional framework (young female, n = 5; mature female, n = 4; young male, n = 4; mature male, n = 5). Because these combined subgroups contained only four to five animals, they were interpreted as supportive analyses rather than as the primary basis for inference. No separate inferential tests were used to compare one sex directly with the other or one age group directly with the other.

Reporter composition within the medial prefrontal cortex was analyzed separately using a one-way repeated-measures ANOVA with reporter category as the within-animal factor, followed by Geisser-Greenhouse correction and Tukey-adjusted multiple comparisons. The pooled cortical cohort included 18 animals; sex- and age-stratified analyses each included 9 animals per group, and age-by-sex groups included 5 young females, 4 mature females, 4 young males, and 5 mature males.

Data are presented as individual-animal values with group means and standard errors of the mean. Statistical significance was defined as p < 0.05, and exact p values were reported where available. For post hoc comparisons, mean differences and 95% confidence intervals were retained to describe the magnitude and precision of observed differences. No statistical tests were applied to reporter-negative DAPI-only cells because these cells were not enumerated by the analysis pipeline. HCR-FISH target-versus-*dapB* overlap percentages were treated descriptively; no inferential statistical comparison between target and negative-control fields was performed.

## Results

### Reporter-defined cellular composition was quantified across forebrain regions of neurological and neuropsychiatric relevance

Reporter-defined cellular composition was quantified in D1-tdTomato/M4-EGFP double-reporter mice across the dorsolateral striatum (DLSTR), dorsomedial striatum (DMSTR), nucleus accumbens (NAc) core, NAc shell, and medial prefrontal cortex (mPFC). The pooled cohort included 18 animals balanced by sex and age. Cells were classified as tdTomato-only, EGFP-only, or tdTomato-EGFP coexpressing, and section-level counts were aggregated within animal before reporter-class composition was calculated. (**Fig. 1**).

Initial 10× survey imaging across neurologically and neuropsychiatrically important forebrain regions revealed localized areas of comparatively high combined tdTomato and EGFP reporter intensity. Threshold-based heatmaps were therefore used to guide higher-magnification sampling within the dorsal striatum, nucleus accumbens, and medial prefrontal cortex, which were subsequently analyzed at the single-cell level. Heatmap intensity itself was not used for quantitative comparisons across animals. (**Fig. 2**).

### Reporter-defined cellular composition differs across striatal subregions

The relative representation of tdTomato-only, EGFP-only, and coexpressing reporter-defined cells differed across dorsal and ventral striatal regions in the pooled cohort (n = 18). Two-way repeated-measures ANOVA identified a significant effect of reporter class (F(1.276, 21.69) = 18.44, p = 0.0001) and a significant region x reporter-class interaction (F(3.780, 64.27) = 22.87, p < 0.0001). The main effect of region was not significant (F(1, 17) = 0.8252, p = 0.3764). Reporter-specific one-way repeated-measures ANOVAs further confirmed significant regional variation in tdTomato-only cells (F(2.634, 44.78) = 36.75, p < 0.0001), EGFP-only cells (F(2.198, 37.36) = 19.41, p < 0.0001), and coexpressing cells (F(2.310, 39.27) = 13.45, p < 0.0001). (**Fig. 5A**).

**Figure 5.**
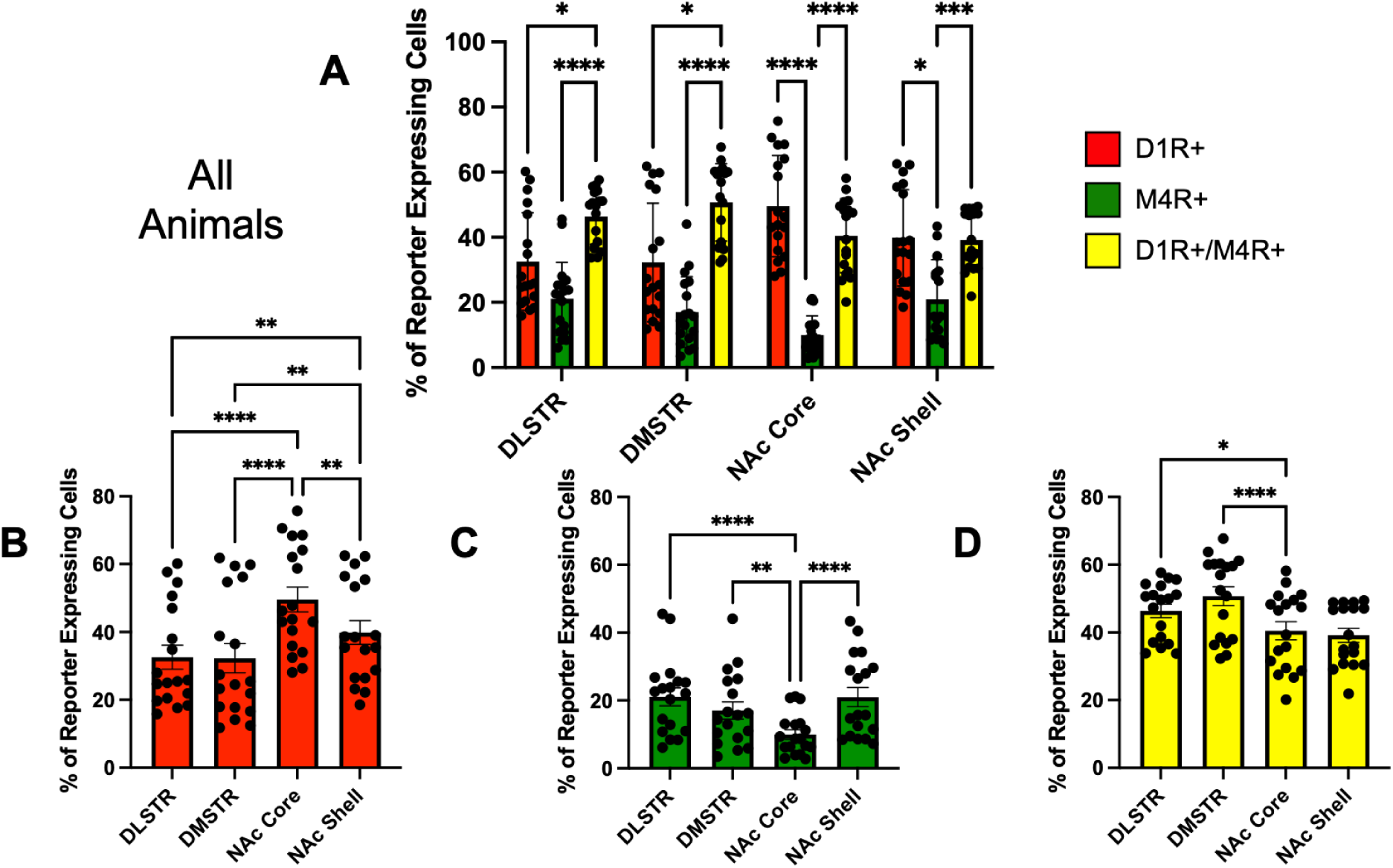
Reporter-defined cellular composition across striatal subregions. **A.** Relative proportions of D1R+ (tdTomato-only), M4R+ (EGFP-only), and D1R+/M4R+ coexpressing cells across the dorsolateral striatum (DLSTR), dorsomedial striatum (DMSTR), nucleus accumbens (NAc) core, and NAc shell in the pooled cohort (n = 18). **B.** Regional distribution of D1R+ cells. **C.** Regional distribution of M4R+ cells. **D.** Regional distribution of D1R+/M4R+ coexpressing cells. Individual points represent animal-level values; bars show group means with error bars. Asterisks indicate significant adjusted pairwise comparisons (*p < 0.05, **p < 0.01, ***p < 0.001, ****p < 0.0001).

In the DLSTR, coexpressing cells represented the largest fraction of the reporter-classified population, comprising 46.36% of cells compared with 32.55% tdTomato-only and 21.09% EGFP-only cells. The coexpressing population was significantly greater than both the tdTomato-only population (p = 0.0375) and the EGFP-only population (p < 0.0001), whereas tdTomato-only and EGFP-only fractions did not differ significantly (p = 0.1581).

A similar coexpression-rich organization was observed in the DMSTR. Coexpressing cells accounted for 50.69% of reporter-classified cells, while tdTomato-only and EGFP-only cells represented 32.27% and 17.04%, respectively. Coexpression exceeded both tdTomato-only (p = 0.0373) and EGFP-only representation (p < 0.0001). The difference between tdTomato-only and EGFP-only populations did not reach significance (p = 0.0752). Thus, both dorsal striatal regions contained substantial reporter coexpression, with the highest mean coexpressing fraction observed in the DMSTR.

The NAc core displayed a distinct composition. tdTomato-only cells comprised 49.57% of the reporter-classified population, compared with 40.50% coexpressing and 9.94% EGFP-only cells. Both tdTomato-only and coexpressing populations were significantly greater than the EGFP-only population (both p < 0.0001), whereas tdTomato-only and coexpressing fractions did not differ significantly (p = 0.3390). These findings indicate that the NAc core retained a substantial coexpressing population but was distinguished by increased tdTomato-only representation and marked depletion of EGFP-only cells.

The NAc shell showed an intermediate organization. tdTomato-only and coexpressing cells represented 39.87% and 39.14% of reporter-classified cells, respectively, and did not differ from one another (p = 0.9885). EGFP-only cells accounted for 20.99% of the population and were significantly less abundant than both tdTomato-only cells (p = 0.0157) and coexpressing cells (p = 0.0003). Thus, unlike the coexpression-dominant dorsal striatum, the accumbens contained similarly represented tdTomato-only and coexpressing populations, with the shift most pronounced in the NAc core.

### Regional comparisons reveal distinct distributions of each reporter-defined population

tdTomato-only representation was similar between DLSTR and DMSTR (32.55% and 32.27%, respectively; p = 0.9963), but was substantially greater in the NAc core (49.57%) than in DLSTR (p < 0.0001), DMSTR (p < 0.0001), or NAc shell (39.87%; p = 0.0013). The NAc shell also contained a greater tdTomato-only fraction than DLSTR (p = 0.0094) and DMSTR (p = 0.0042). (**Fig. 5B**).

EGFP-only representation was 21.09% in DLSTR, 17.04% in DMSTR, 9.94% in NAc core, and 20.99% in NAc shell. DLSTR exceeded DMSTR (p = 0.0085) and NAc core (p < 0.0001), but not NAc shell (p > 0.9999). DMSTR exceeded NAc core (p = 0.0024) but did not differ from NAc shell (p = 0.0746), while NAc core was lower than NAc shell (p < 0.0001). (**Fig. 5C**).

Coexpression was highest in DMSTR (50.69%), exceeding DLSTR (46.36%; p = 0.0262), NAc core (40.50%; p < 0.0001), and NAc shell (39.14%; p = 0.0006). DLSTR coexpression was also greater than NAc core (p = 0.0317) and NAc shell (p = 0.0233), whereas NAc core and NAc shell did not differ (p = 0.9526). (**Fig. 5D**).

### Striatal regional organization is broadly preserved across sex and age groups

The region-dependent organization of reporter-defined populations was not restricted to a single age group (**Fig. 6**). Region x reporter interactions were significant in young animals (**Fig. 6A-D**; p = 0.0001) and mature animals (**Fig. 6E-H**; p < 0.0001).

**Figure 6.**
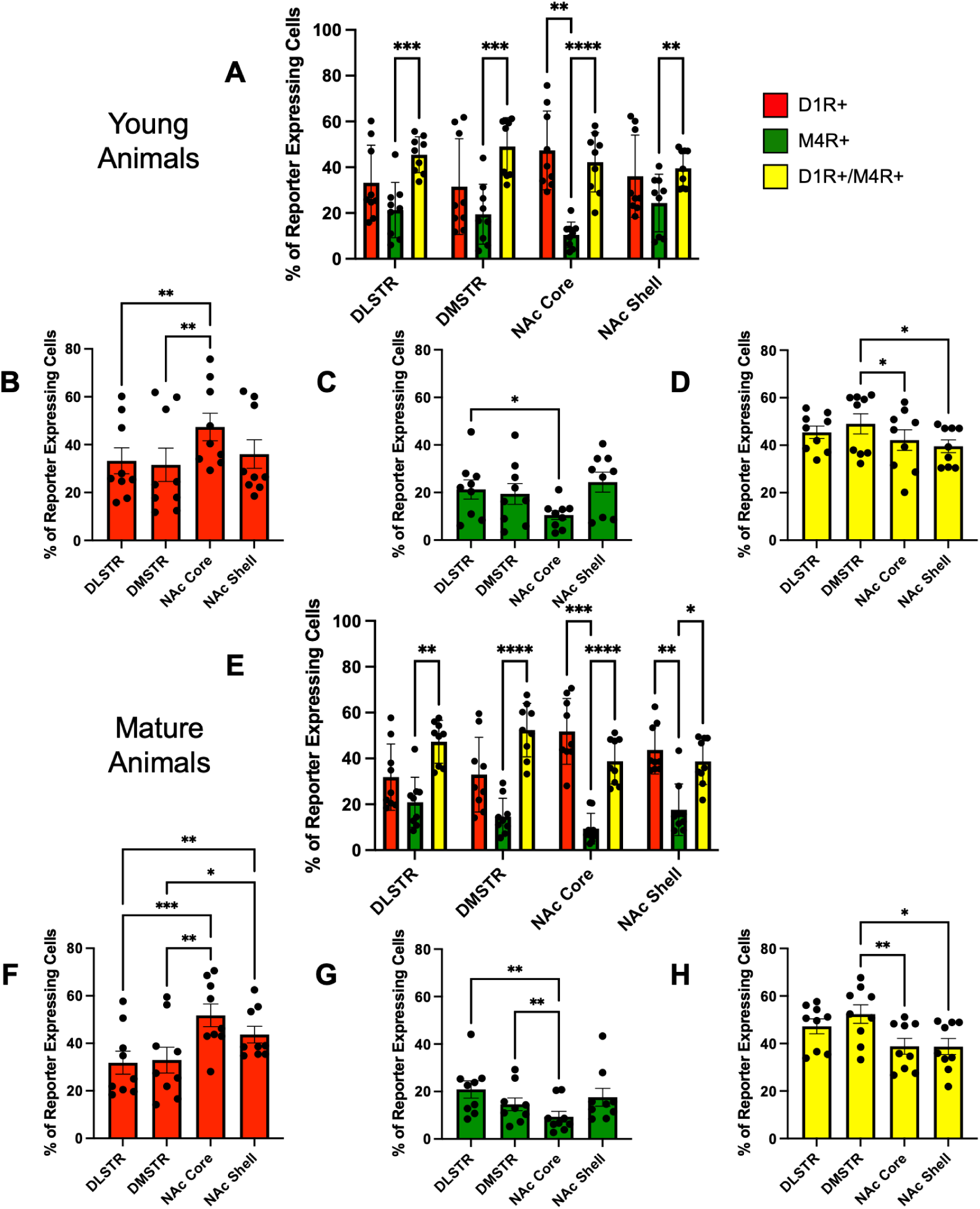
Age-stratified reporter-defined cellular composition across striatal subregions. **A.** Relative proportions of D1R+ (tdTomato-only), M4R+ (EGFP-only), and D1R+/M4R+ coexpressing cells across the dorsolateral striatum (DLSTR), dorsomedial striatum (DMSTR), nucleus accumbens (NAc) core, and NAc shell in young animals. **B.** Regional distribution of D1R+ cells in young animals. **C.** Regional distribution of M4R+ cells in young animals. **D.** Regional distribution of D1R+/M4R+ coexpressing cells in young animals. **E.** Relative proportions of D1R+, M4R+, and D1R+/M4R+ coexpressing cells across the same striatal subregions in mature animals. **F.** Regional distribution of D1R+ cells in mature animals. **G.** Regional distribution of M4R+ cells in mature animals. **H.** Regional distribution of D1R+/M4R+ coexpressing cells in mature animals. Individual points represent animal-level values; bars show group means with error bars. Asterisks indicate significant adjusted pairwise comparisons (*p < 0.05, **p < 0.01, ***p < 0.001, ****p < 0.0001).

Reporter-specific regional effects were significant in young animals for tdTomato-only (F(2.837, 22.70) = 16.42, p < 0.0001), EGFP-only (F(1.866, 14.93) = 11.31, p = 0.0012), and coexpressing cells (F(2.266, 18.13) = 5.146, p = 0.0143), and in mature animals for tdTomato-only (F(2.225, 17.80) = 24.59, p < 0.0001), EGFP-only (F(1.466, 11.73) = 10.41, p = 0.0042), and coexpressing cells (F(2.030, 16.24) = 8.549, p = 0.0028).

The same overall organization was retained when animals were grouped by sex (**Fig. 7**). The region x reporter interaction remained significant in females (**Fig. 7A-D**; p = 0.0004) and males (**Fig. 7E-H**; p < 0.0001). Reporter-specific regional effects were also significant in females for tdTomato-only (F(2.474, 19.80) = 10.73, p = 0.0004), EGFP-only (F(1.501, 12.01) = 14.15, p = 0.0012), and coexpressing cells (F(2.297, 18.37) = 6.082, p = 0.0075), and in males for tdTomato-only (F(2.060, 16.48) = 31.18, p < 0.0001), EGFP-only (F(2.003, 16.02) = 10.19, p = 0.0014), and coexpressing cells (F(2.150, 17.20) = 8.357, p = 0.0025).

**Figure 7.**
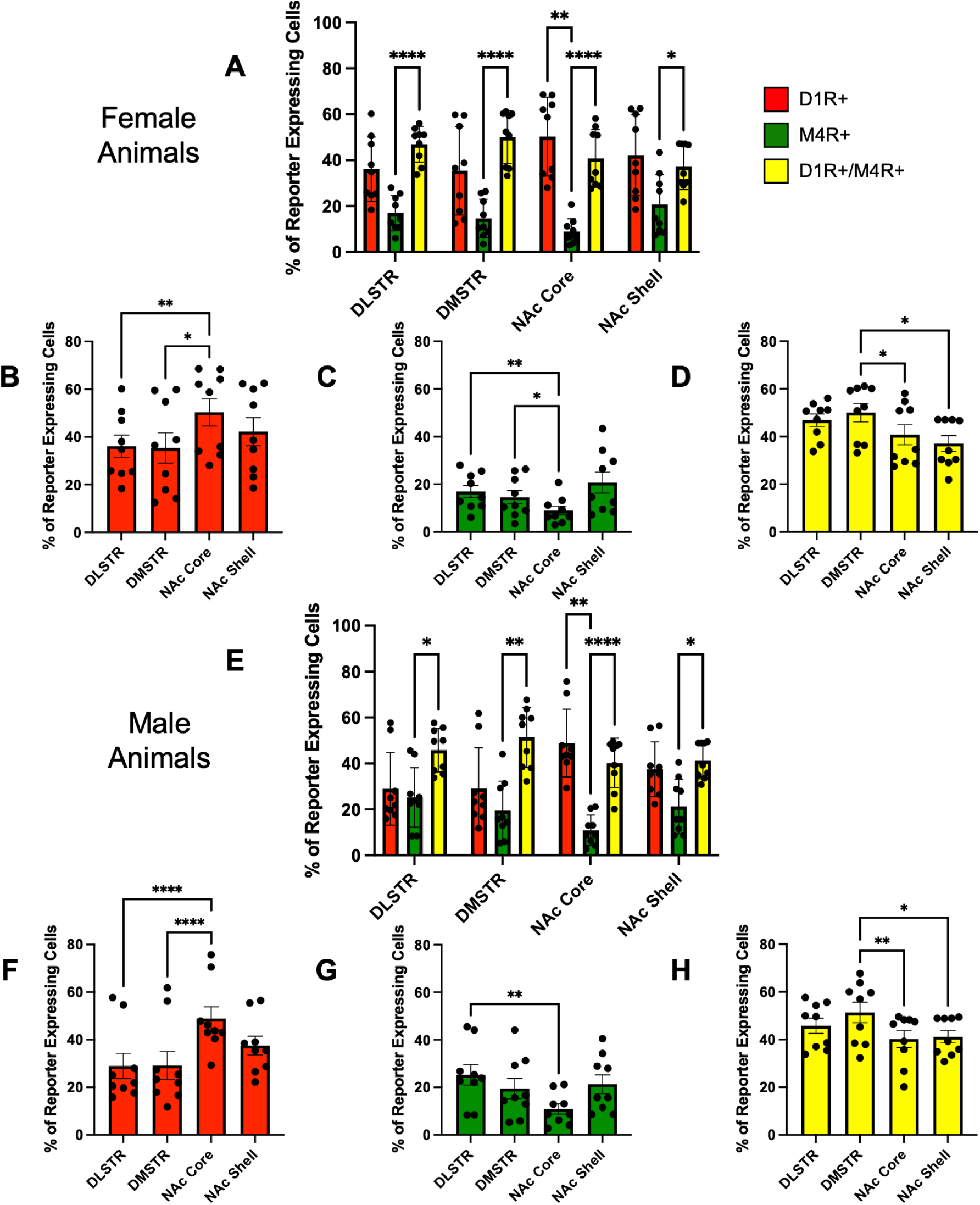
Sex-stratified reporter-defined cellular composition across striatal subregions. **A.** Relative proportions of D1R+ (tdTomato-only), M4R+ (EGFP-only), and D1R+/M4R+ coexpressing cells across the dorsolateral striatum (DLSTR), dorsomedial striatum (DMSTR), nucleus accumbens (NAc) core, and NAc shell in female animals. **B.** Regional distribution of D1R+ cells in female animals. **C.** Regional distribution of M4R+ cells in female animals. **D.** Regional distribution of D1R+/M4R+ coexpressing cells in female animals. **E.** Relative proportions of D1R+, M4R+, and D1R+/M4R+ coexpressing cells across the same striatal subregions in male animals. **F.** Regional distribution of D1R+ cells in male animals. **G.** Regional distribution of M4R+ cells in male animals. **H.** Regional distribution of D1R+/M4R+ coexpressing cells in male animals. Individual points represent animal-level values; bars show group means with error bars. Asterisks indicate significant adjusted pairwise comparisons (*p < 0.05, **p < 0.01, ***p < 0.001, ****p < 0.0001).

Further subdivision by both age and sex produced groups of four to five animals and was therefore interpreted supportively. Young females (**Supplementary Fig. 1A-D**) showed significant reporter-specific regional effects for tdTomato-only (F(1.368, 5.474) = 7.673, p = 0.0310), EGFP-only (F(1.247, 4.989) = 10.34, p = 0.0212), and coexpressing cells (F(1.617, 6.467) = 6.344, p = 0.0335). No reporter-specific regional omnibus reached significance in mature females (**Supplementary Fig. 2A-D**) (tdTomato-only p = 0.0617; EGFP-only p = 0.1441; coexpressing p = 0.1389). In young males (**Supplementary Fig. 1E-H**), only the tdTomato-only regional effect was significant (F(2.282, 6.847) = 9.445, p = 0.0097), whereas EGFP-only (p = 0.1291) and coexpressing (p = 0.3271) effects were not. Mature males (**Supplementary Fig. 2E-H**) showed significant regional effects for tdTomato-only (F(1.297, 5.188) = 27.32, p = 0.0025), EGFP-only (F(1.170, 4.679) = 21.17, p = 0.0062), and coexpressing cells (F(1.796, 7.186) = 19.83, p = 0.0013).

### Medial prefrontal cortex exhibits an EGFP-only-predominant reporter composition

Reporter-defined cellular organization in the sampled medial prefrontal cortex differed markedly from that observed in the striatum. Across the pooled cohort (n = 18), EGFP-only cells represented 61.01% of reporter-classified cells, whereas tdTomato-only and coexpressing cells accounted for 20.03% and 18.95%, respectively. One-way repeated-measures ANOVA identified a significant effect of reporter class (F(1.546, 26.28) = 58.35, p < 0.0001). EGFP-only representation was greater than both tdTomato-only (p < 0.0001) and coexpressing representation (p < 0.0001), whereas tdTomato-only and coexpressing fractions did not differ (p = 0.9358). (**Fig. 8A**).

**Figure 8.**
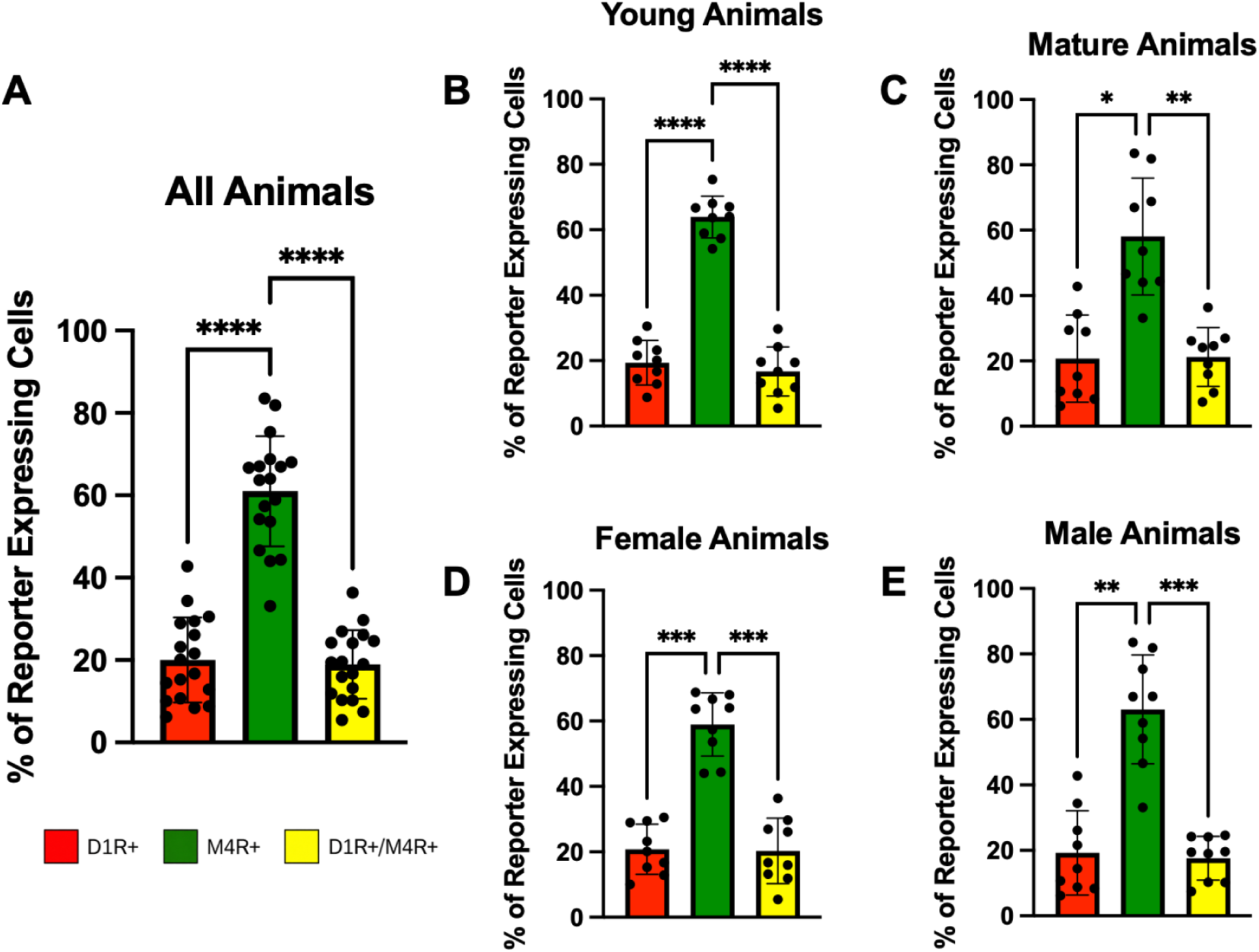
Reporter-defined cellular composition in the medial prefrontal cortex across the pooled cohort and major demographic subgroups. **A.** Relative proportions of D1R+ (tdTomato-only), M4R+ (EGFP-only), and D1R+/M4R+ coexpressing cells in the pooled cohort. **B.** Reporter-defined cellular composition in young animals. **C.** Reporter-defined cellular composition in mature animals. **D.** Reporter-defined cellular composition in female animals. **E.** Reporter-defined cellular composition in male animals. Individual points represent animal-level values; bars show group means with error bars. Asterisks indicate significant adjusted pairwise comparisons (*p < 0.05, **p < 0.01, ***p < 0.001, ****p < 0.0001).

This EGFP-only predominance was evident in both sexes. In females (**Fig. 8D**; n = 9), tdTomato-only, EGFP-only, and coexpressing cells represented 20.79%, 58.95%, and 20.26%, respectively; reporter class differed significantly overall (p < 0.0001), with EGFP-only representation exceeding tdTomato-only (p = 0.0001) and coexpressing populations (p = 0.0005). In males (**Fig. 8E**; n = 9), the corresponding fractions were 19.27%, 63.07%, and 17.65%; reporter class again differed significantly (p = 0.0005), with EGFP-only cells exceeding tdTomato-only (p = 0.0048) and coexpressing cells (p = 0.0006).

Cortical reporter composition was similarly stable across age groups. Young animals (**Fig. 8B**; n = 9) showed 19.37% tdTomato-only, 63.92% EGFP-only, and 16.70% coexpressing cells; reporter class differed significantly (p < 0.0001), with EGFP-only representation exceeding both other populations (both p < 0.0001). Mature animals (**Fig. 8C**; n = 9) showed 20.69% tdTomato-only, 58.10% EGFP-only, and 21.21% coexpressing cells; the reporter-class effect remained significant (p = 0.0021), with EGFP-only cells exceeding tdTomato-only (p = 0.0146) and coexpressing cells (p = 0.0055).

### Cortical EGFP predominance is retained across age-by-sex subgroups

The same qualitative cortical organization was observed when animals were subdivided simultaneously by age and sex. Young females (**Supplementary Fig. 3A**; n = 5) showed 20.67% tdTomato-only, 63.95% EGFP-only, and 15.38% coexpressing cells (overall p = 0.0007; EGFP-only vs tdTomato-only p = 0.0003; EGFP-only vs coexpressing p = 0.0020). Mature females (**Supplementary Fig. 3C**; n = 4) showed 20.95%, 52.70%, and 26.36%, respectively (overall p = 0.0290), although no individual Tukey comparison reached significance. Young males (**Supplementary Fig. 3B**; n = 4) showed 17.75%, 63.89%, and 18.36% (overall p = 0.0021; EGFP-only vs tdTomato-only p = 0.0209; EGFP-only vs coexpressing p = 0.0136). Mature males (**Supplementary Fig. 3D**; n = 5) showed 20.49%, 62.42%, and 17.09% (overall p = 0.0429); no Tukey comparison reached significance, with EGFP-only versus coexpressing approaching the threshold (p = 0.0507).

### Reporter-defined organization differs fundamentally between striatum and cortex

Taken together, the striatal and cortical analyses revealed region-specific reporter organization rather than a uniform relationship between tdTomato and EGFP expression across forebrain regions. Dorsal striatum contained a large coexpressing population, with coexpression maximal in the DMSTR. The nucleus accumbens shifted toward increased tdTomato-only representation and reduced EGFP-only representation, particularly in the NAc core. In contrast, the medial prefrontal cortex was dominated by EGFP-only cells, with tdTomato-only and coexpressing populations each representing approximately one-fifth of reporter-classified cells. These regional patterns were broadly preserved across sex and age.

### HCR-FISH supports correspondence between endogenous receptor transcripts and reporter fluorescence

HCR-FISH was used to assess correspondence between endogenous Drd1 and Chrm4 transcript signal and the respective tdTomato and EGFP reporters. In sections hybridized with the Drd1 target probe (**Fig. 9A**), 95.4% of detected HCR puncta overlapped tdTomato-positive signal, compared with 58.8% apparent overlap in dapB negative-control sections. Representative images showed concentrated target-probe HCR signal associated with tdTomato-positive somata, whereas the negative-control signal lacked the same reporter-associated distribution.

**Figure 9.**
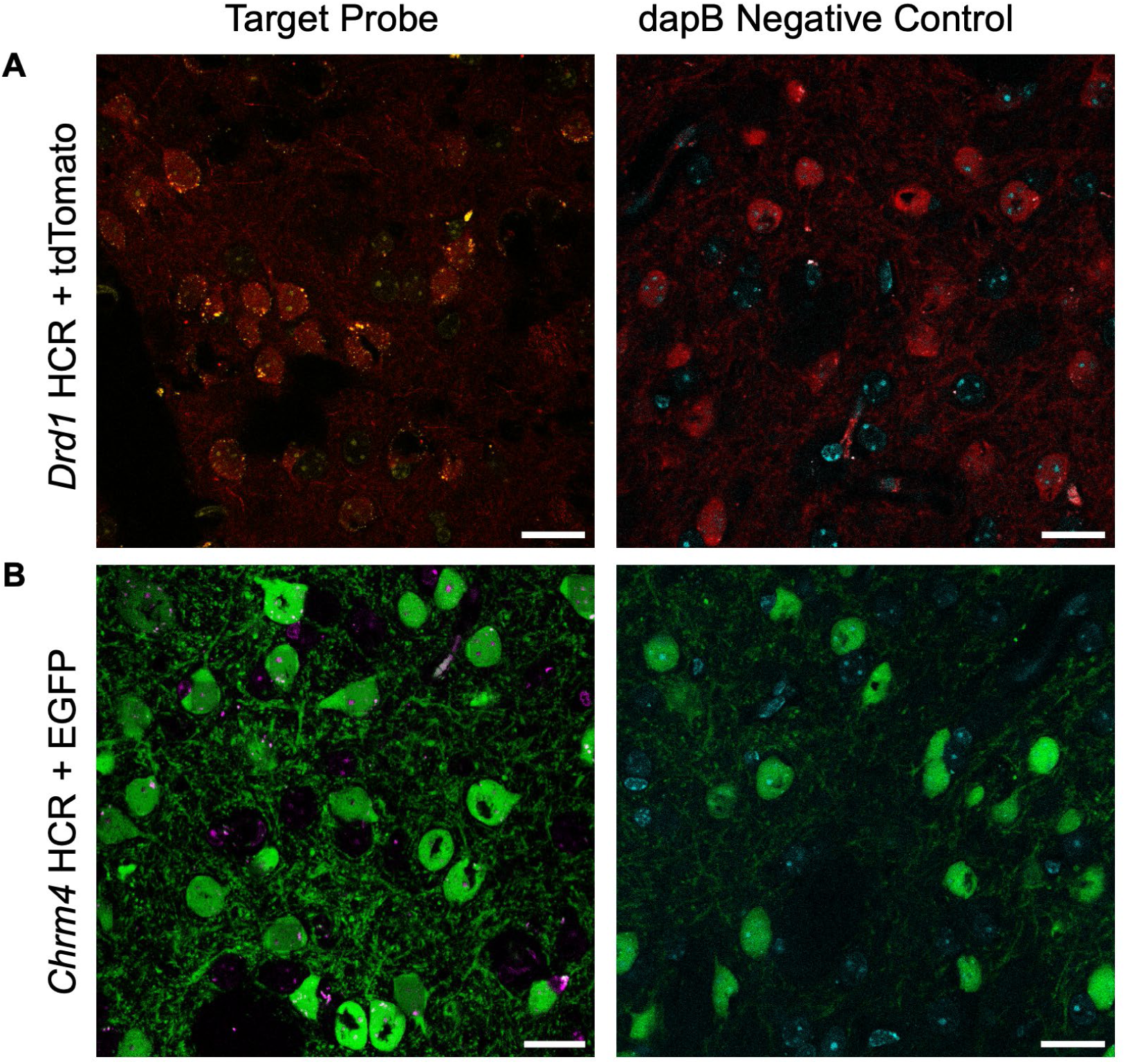
HCR-FISH validation of *Drd1-*tdTomato and *Chrm4-*EGFP reporter expression. **A.** Representative Drd1 HCR-FISH signal in tdTomato reporter tissue (left) compared with the dapB negative control (right). Pooled HCR puncta overlap with tdTomato-positive signal was 95.4% for Drd1 and 58.8% for *dapB*. **B.** Representative Chrm4 HCR-FISH signal in EGFP reporter tissue (left) compared with *dapB* (right). Pooled HCR puncta overlap with EGFP-positive signal was 77.6% for Chrm4 and 52.2% for *dapB*. HCR signal was acquired in the Alexa Fluor 647 channel and pseudocolored yellow for *Drd1*, magenta for *Chrm4*, and cyan for *dapB*; tdTomato and EGFP reporter fluorescence are shown in red and green, respectively. Scale bars, 20 µm.

A similar relationship was observed for Chrm4 and the M4-EGFP reporter (**Fig. 9B**). In target-probe sections, 77.6% of detected HCR puncta overlapped EGFP-positive signal, compared with 52.2% in dapB negative-control sections. Together, these findings support spatial correspondence between endogenous Drd1 and Chrm4 transcript localization and expression of their respective fluorescent reporters (i.e., tdTomato and EGFP).

## Discussion

The present findings establish a regionally heterogeneous relationship between D1R- and M4R-associated cellular populations across the cortex and striatum. Reporter coexpression was substantial throughout the dorsal and ventral striatum, but the relative composition of tdTomato-only, EGFP-only, and coexpressing populations differed markedly across anatomical regions. Coexpression was particularly prominent in the dorsal striatum and was greatest in the dorsomedial striatum, whereas the nucleus accumbens shifted toward a larger tdTomato-only population and reduced EGFP-only representation, most clearly in the NAc core. The sampled medial prefrontal cortex showed the opposite organization, with EGFP-only cells constituting the predominant reporter-defined population. These broader regional patterns were retained across sex and age, indicating that the organization observed in the pooled cohort was not driven by a single demographic subset. HCR-FISH further supported correspondence between endogenous *Drd1* and *Chrm4* transcript localization and the respective tdTomato and EGFP reporters. Together, these findings indicate that the cellular opportunity for M4R-mediated modulation of D1R-associated neurons is substantial but varies considerably with anatomical context.

The widespread coexpressing population observed throughout the striatum provides anatomical support for considering M4R as a receptor-context target within D1R-associated neuronal populations. M4R is known to be coexpressed with D1R in a subset of striatonigral neurons, and selective deletion of M4R from D1R-expressing neurons alters dopamine-dependent behavioral responses.^17,19^ In each of the four striatal and accumbal regions examined here, cells carrying both reporters represented a substantial fraction of the reporter-classified population, indicating that D1R- and M4R-associated expression frequently occurs within the same cells rather than exclusively in separate populations occupying the same tissue. This distinction is pharmacologically important because modulation through M4R could alter the signaling environment of D1R-associated neurons from within the same cellular population rather than acting only indirectly through neighboring cells or parallel circuit elements.^19^ Reporter coexpression, however, does not establish equivalent receptor abundance, subcellular colocalization, or functional coupling within every coexpressing cell. The present measurements therefore define the regional cellular opportunity for D1R-M4R integration rather than the magnitude or mechanism of that interaction.

The particularly high relative coexpression observed in the DMSTR identifies this region as the strongest anatomical example of this receptor-context relationship. Coexpressing cells accounted for approximately half of the reporter-defined population and were more prevalent in the DMSTR than in the DLSTR, NAc core, or NAc shell. This regional distinction is notable because the dorsolateral and dorsomedial striatum participate in partially distinct functional circuits. The DLSTR has been broadly associated with sensorimotor and habit-related control, whereas the DMSTR contributes more prominently to associative and goal-directed behavior.^37,38^ Greater relative reporter coexpression in the DMSTR therefore suggests that M4R may be positioned within a particularly large D1R-associated cellular population in this region. Such an arrangement could provide a means of influencing D1R-associated signaling without directly targeting D1R itself, an idea that is relevant to continuing interest in M4R-selective pharmacology as a strategy for modifying dopamine-linked function.^20–25,27–31^ The present analysis does not establish that M4R signaling is stronger in the DMSTR, however, because the measurements describe relative population composition rather than receptor abundance, signaling capacity, or absolute cellular density.

The nucleus accumbens provided an important contrast to the dorsal striatum. In the NAc core, tdTomato-only cells constituted approximately half of the reporter-classified population, EGFP-only cells were markedly reduced, and coexpressing cells nevertheless remained a substantial component. The NAc shell showed a related but less pronounced distribution, with tdTomato-only and coexpressing cells present at similar proportions and EGFP-only cells comprising the smaller population. The dorsal-to-ventral transition was therefore characterized less by a loss of coexpression than by redistribution of the two single-positive populations. This distinction is relevant because the NAc core and shell differ in connectivity, dopaminergic innervation, and contributions to motivated behavior.^39,40^ The core is more strongly associated with cue-guided reward seeking, whereas the shell contributes more prominently to contextual regulation of reward-related behavior.^41–43^ More broadly, the nucleus accumbens integrates cognitive and affective information involved in motivated behavior and action selection.^53^ The present findings indicate that the cellular relationship between D1R- and M4R-associated populations differs even between closely related accumbal compartments and should not be assumed to be uniform across ventral striatum.

The prominence of the tdTomato-only population in the NAc core may also define an important boundary to an M4R-directed receptor-context strategy. A substantial fraction of D1R-associated reporter cells in this region lacked detectable M4-EGFP signal and may therefore fall outside the direct cellular reach of an M4R-directed intervention, even though coexpression remained considerable. At the same time, the relatively small EGFP-only population suggests that M4R-associated reporter expression in the core was concentrated more strongly within cells that also carried the D1R reporter than within a separate M4R-associated population. The core-shell redistribution of these reporter classes therefore indicates that the potential cellular selectivity of M4R pharmacology can differ even within the nucleus accumbens. One possibility is that tdTomato-only and coexpressing D1R-associated cells represent populations with different projection targets, local connectivity, or signaling machinery. The present data cannot resolve those cellular identities directly, but they provide an anatomical basis for testing whether M4R-directed pharmacology preferentially influences specific D1R-associated circuits rather than acting uniformly across the full D1R-associated population.

The sampled medial prefrontal cortical region showed a clearly different organization. EGFP-only cells constituted approximately three-fifths of reporter-classified cells, whereas tdTomato-only and coexpressing populations were each present at substantially lower proportions. Thus, within this cortical region, M4R-associated reporter expression was more often detected without corresponding D1R-associated reporter expression than within the overlapping populations prominent in the dorsal striatum. This regional difference suggests that the relationship between M4R pharmacology and D1R-associated cellular context is not uniform across the brain. An M4R-directed intervention may be closely aligned with D1R-associated populations in regions with extensive reporter coexpression while engaging a compositionally different set of reporter-defined cells in cortex. The cortical result also argues that the extensive striatal coexpression was not simply an inevitable consequence of combining the two reporter lines, but instead reflected regionally distinct cellular organization. The medial prefrontal cortex integrates contextual and mnemonic information involved in decision-making and adaptive behavior,^54^ providing a functionally distinct circuit in which to observe this contrasting arrangement. Because the cortical ROI combined portions of the anterior cingulate, prelimbic, and infralimbic cortices, the present analysis cannot determine whether this composition was shared uniformly across those subdivisions. Finer anatomical sampling may therefore reveal local cortical populations with greater or lesser D1R-M4R reporter overlap than was apparent in the combined ROI.

The principal regional organization was broadly retained when the cohort was divided by sex or age. Coexpression remained prominent within the dorsal striatum in female, male, young, and mature animals, while the NAc core repeatedly showed increased tdTomato-only representation and reduced EGFP-only representation. The recurrence of these features indicates that the pooled organization was not driven by a single subgroup. The tdTomato-rich organization of the NAc core was also evident across multiple demographic groups rather than being restricted to a particular age-by-sex combination, supporting its interpretation as a broader anatomical feature of this region. Combined age-by-sex groups were smaller and showed correspondingly greater variability, so differences in statistical significance among those groups should not themselves be interpreted as evidence for age- or sex-dependent organization. These subgroup analyses instead support the overall stability of the regional pattern while leaving open the possibility of subtler demographic effects that would require a study specifically powered to test sex-by-age interactions.

HCR-FISH strengthened the interpretation of the fluorescent reporter populations as anatomically meaningful proxies for endogenous receptor-associated expression. *Drd1* target-probe signal showed strong spatial correspondence with tdTomato fluorescence, with 95.4% of detected puncta overlapping reporter-positive signal, compared with 58.8% apparent overlap under the dapB negative-control condition. Similarly, 77.6% of *Chrm4* puncta overlapped EGFP-positive signal, compared with 52.2% in the corresponding dapB comparison. These findings provide transcript-level support for both reporter systems and increase confidence that the regional reporter distributions identify populations with endogenous capacity to express the corresponding receptors. The validation should nevertheless be interpreted as an assessment of spatial correspondence rather than a quantitative measure of transcript abundance. HCR-FISH was performed as a validation experiment in a single animal, and target and negative-control conditions were not subjected to an inferential statistical comparison. Reporter-positive fluorescence also occupied a substantial portion of the analyzed fields, such that some punctum overlap would be expected from spatial proximity alone. The HCR data therefore support correspondence between endogenous transcripts and reporter-defined populations but do not establish receptor protein abundance, surface localization, or functional signaling competence.

Several limitations of the reporter-based approach should also be considered. BAC reporter fluorescence provides an indirect readout of receptor-associated cellular identity and does not measure endogenous receptor protein itself. Reporter expression may differ from receptor protein as a consequence of transcriptional regulation, fluorophore stability, or properties of the transgenic constructs. Direct protein-level mapping of GPCR expression is itself complicated by limitations in the specificity and reliability of many GPCR antibodies, providing additional rationale for combining reporter- and transcript-based approaches.^52^ The present analysis also quantified the relative proportions of tdTomato-only, EGFP-only, and coexpressing cells among reporter-classified cells. These values therefore should not be interpreted as the fraction of all cells within a region expressing either receptor or as measures of absolute cellular density. Similarly, fluorescence intensity was not used for quantitative comparisons of receptor abundance across animals. Finally, the automated classification pipeline necessarily relies on operational definitions of reporter positivity and coexpression. Standardized imaging, DAPI-gated cell identification, consistent classification criteria, and aggregation of section-level data within animal were used to reduce technical variability, but the resulting populations remain reporter-defined anatomical categories rather than direct measurements of receptor stoichiometry.

Taken together, the present study establishes a clear regional organization of D1R- and M4R-associated cellular populations across the cortex and striatum. Dorsal striatum contained a substantial coexpressing population, with the greatest relative coexpression in the DMSTR. The nucleus accumbens shifted toward greater tdTomato-only and reduced EGFP-only representation, particularly in the NAc core, while the medial prefrontal cortex showed a markedly different EGFP-only-predominant organization. These patterns were broadly retained across the sexes and ages examined and were supported by transcript-level validation of both reporter systems. The findings therefore argue against a single forebrain-wide relationship between D1R- and M4R-associated populations. Instead, they identify anatomical context as a major determinant of their cellular relationship and provide a regional framework for future studies examining how M4R-directed pharmacology influences D1R-associated signaling within distinct corticostriatal circuits.

## Acknowledgements

This work is supported by Northeastern University Tier 1 grant (H.Y. and J.R.M.) and Northeastern University Startup Funds (H.Y.).

## Author contributions

**Rian Garland**: Data curation, Formal analysis, Investigation, Visualization, Writing, **Maya Dickson**: Data curation, Investigation, **Alice Lee**: Data curation, Investigation, **Ali Noorouzi**: Data curation, Investigation, **Hayato Umaoka**: Data curation, Investigation, **Ana Semeano**: Investigation, **James Monaghan**: Data curation, Formal analysis, Funding acquisition, Resources, Supervision, **Hideaki Yano**: Conceptualization, Data curation, Formal analysis, Funding acquisition, Resources, Supervision, Writing

## Supplementary Figures

**Supplementary Figure 1.**
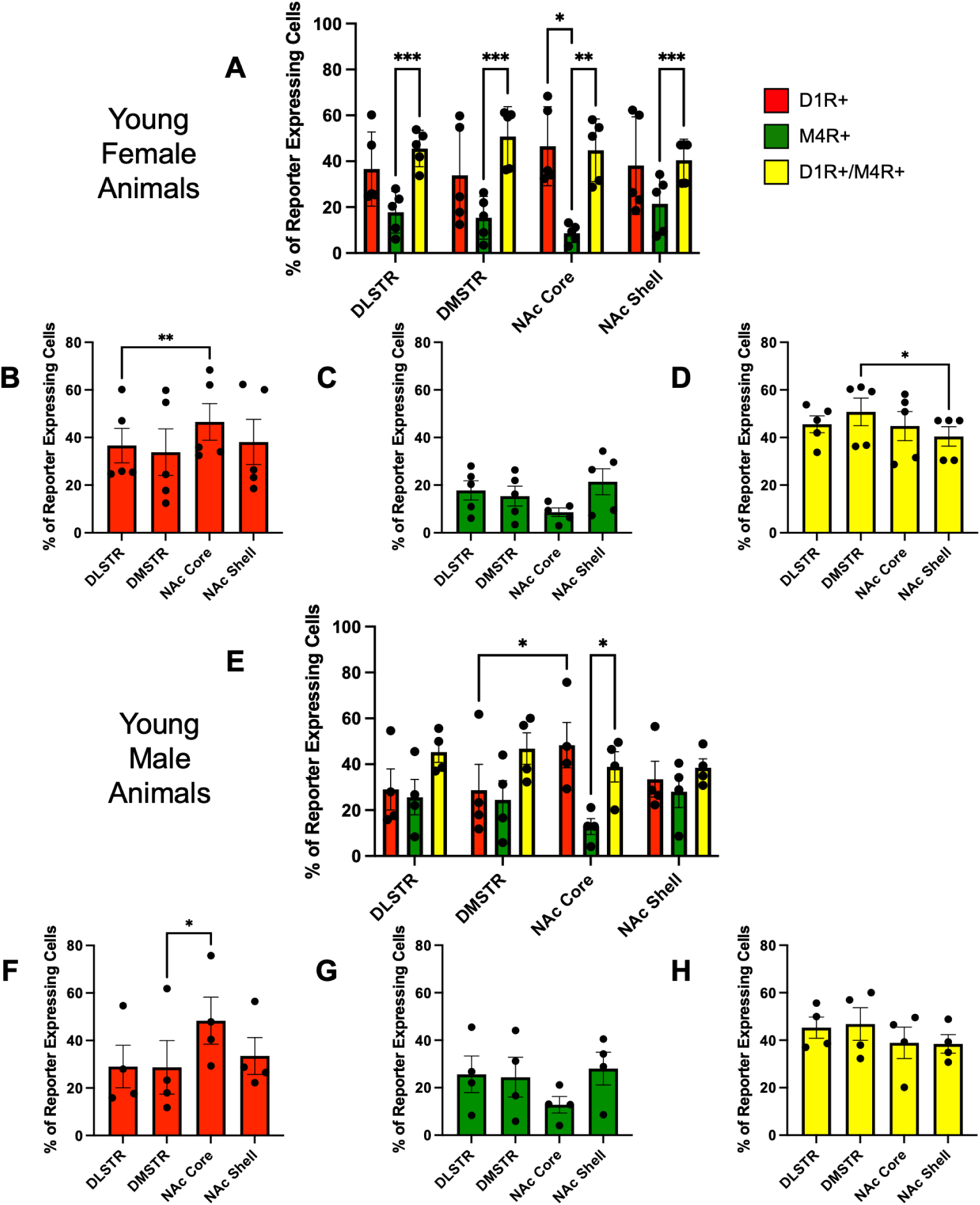
Reporter-defined cellular composition across striatal subregions in young female and young male animals. **A.** Relative proportions of D1R+ (tdTomato-only), M4R+ (EGFP-only), and D1R+/M4R+ coexpressing cells across the dorsolateral striatum (DLSTR), dorsomedial striatum (DMSTR), nucleus accumbens (NAc) core, and NAc shell in young female animals. **B.** Regional distribution of D1R+ cells in young female animals. **C.** Regional distribution of M4R+ cells in young female animals. **D.** Regional distribution of D1R+/M4R+ coexpressing cells in young female animals. **E.** Relative proportions of D1R+, M4R+, and D1R+/M4R+ coexpressing cells across the same striatal subregions in young male animals. **F.** Regional distribution of D1R+ cells in young male animals. **G.** Regional distribution of M4R+ cells in young male animals. **H.** Regional distribution of D1R+/M4R+ coexpressing cells in young male animals. Individual points represent animal-level values; bars show group means with error bars. Asterisks indicate significant adjusted pairwise comparisons (*p < 0.05, **p < 0.01, ***p < 0.001, ****p < 0.0001).

**Supplementary Figure 2.**
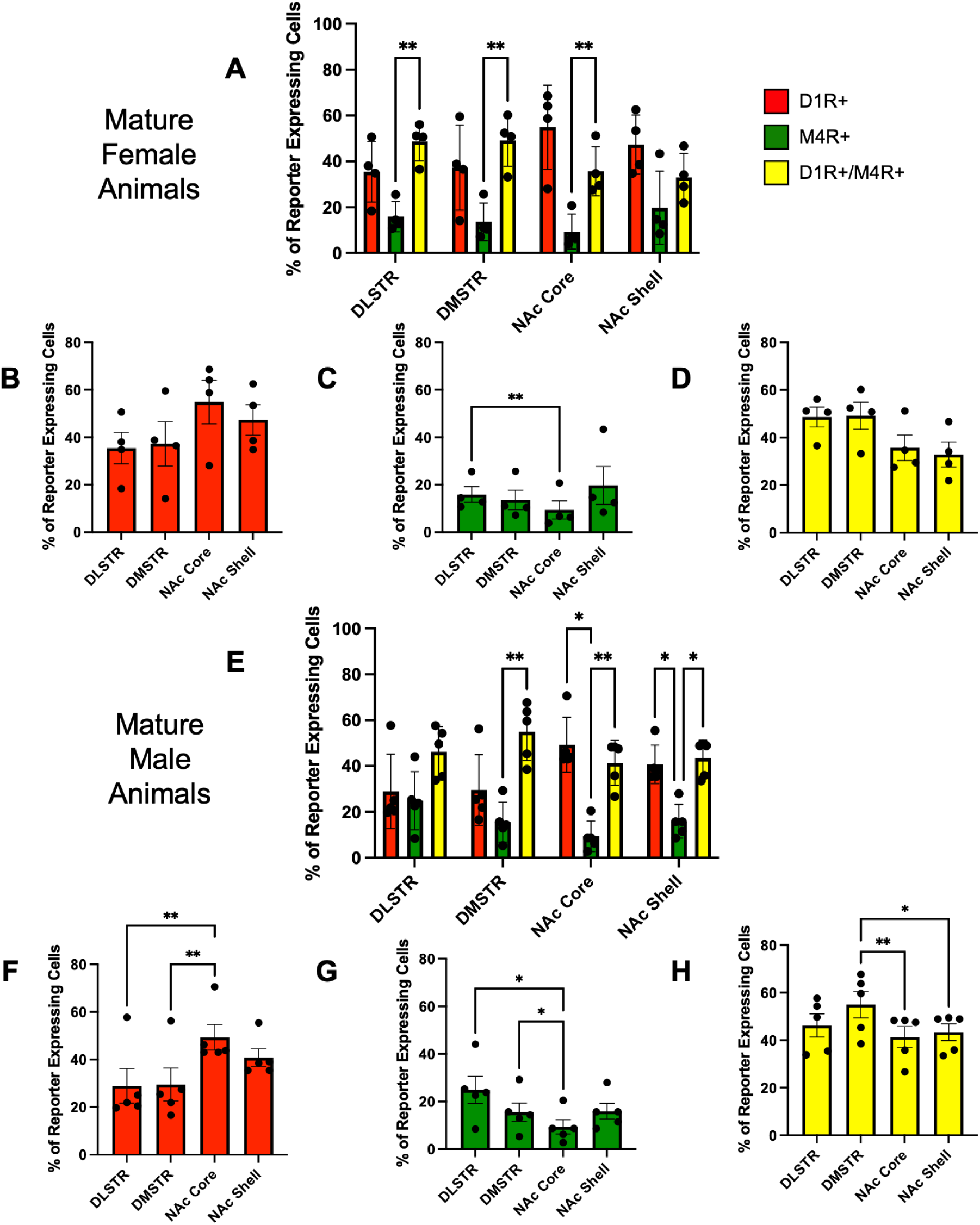
Reporter-defined cellular composition across striatal subregions in mature female and mature male animals. **A.** Relative proportions of D1R+ (tdTomato-only), M4R+ (EGFP-only), and D1R+/M4R+ coexpressing cells across the dorsolateral striatum (DLSTR), dorsomedial striatum (DMSTR), nucleus accumbens (NAc) core, and NAc shell in mature female animals. **B.** Regional distribution of D1R+ cells in mature female animals. **C.** Regional distribution of M4R+ cells in mature female animals. **D.** Regional distribution of D1R+/M4R+ coexpressing cells in mature female animals. **E.** Relative proportions of D1R+, M4R+, and D1R+/M4R+ coexpressing cells across the same striatal subregions in mature male animals. **F.** Regional distribution of D1R+ cells in mature male animals. **G.** Regional distribution of M4R+ cells in mature male animals. **H.** Regional distribution of D1R+/M4R+ coexpressing cells in mature male animals. Individual points represent animal-level values; bars show group means with error bars. Asterisks indicate significant adjusted pairwise comparisons (*p < 0.05, **p < 0.01, ***p < 0.001, ****p < 0.0001).

**Supplementary Figure 3.**
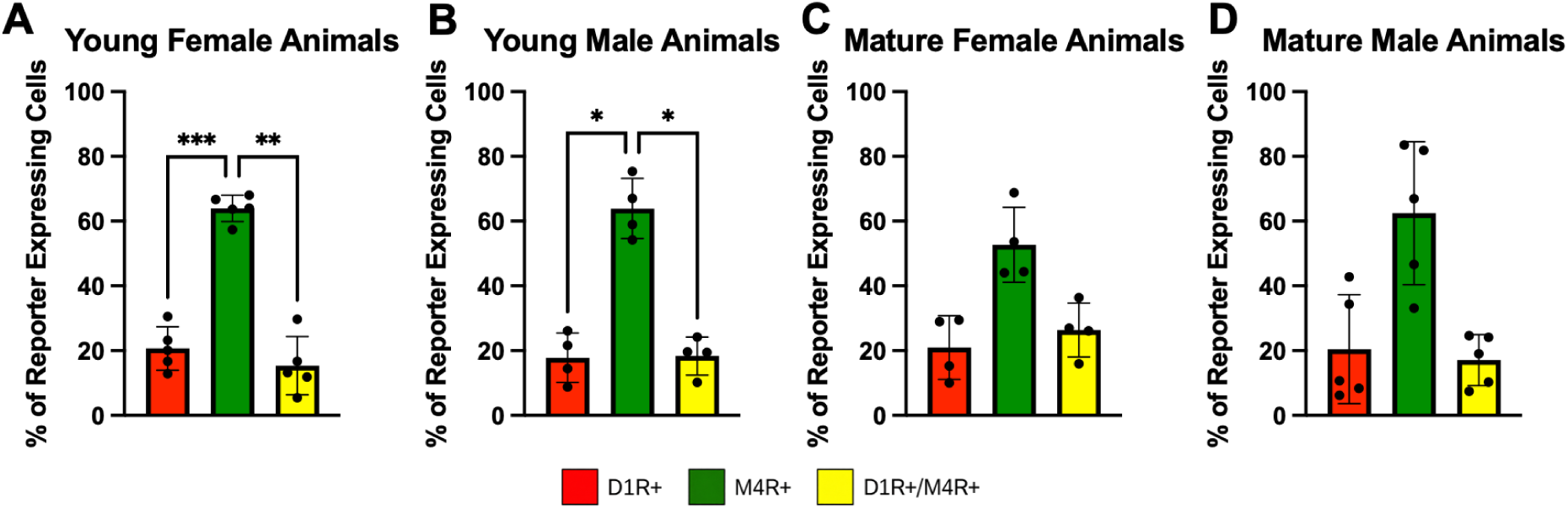
Reporter-defined cellular composition in the medial prefrontal cortex across age-by-sex subgroups. **A.** Relative proportions of D1R+ (tdTomato-only), M4R+ (EGFP-only), and D1R+/M4R+ coexpressing cells in young female animals. **B.** Reporter-defined cellular composition in young male animals. **C.** Reporter-defined cellular composition in mature female animals. **D.** Reporter-defined cellular composition in mature male animals. Individual points represent animal-level values; bars show group means with error bars. Asterisks indicate significant adjusted pairwise comparisons (*p < 0.05, **p < 0.01, ***p < 0.001, ****p < 0.0001).

